# Hidden Talents: Poly (I:C)-induced maternal immune activation improves mouse visual discrimination performance and reversal learning in a sex-dependent manner

**DOI:** 10.1101/2021.02.15.431275

**Authors:** Xin Zhao, Hieu Tran, Holly DeRosa, Ryland C. Roderick, Amanda C. Kentner

## Abstract

While there is a strong focus on the negative consequences of maternal immune activation (MIA) on developing brains, very little attention is directed towards potential advantages of early life challenges. In this study we utilized a polyinosine-polycytidylic acid (poly(I:C)) MIA model to test visual discrimination (VD) and reversal learning (RL) in mice using touchscreen technology. Significant sex differences emerged in that MIA reduced the latency for males to make a correct choice in the VD task while females reached criterion sooner, made fewer errors, and utilized fewer correction trials in RL compared to saline controls. These surprising improvements were accompanied by the sex-specific upregulation of several genes critical to cognitive functioning, indicative of compensatory plasticity in response to MIA. In contrast, when exposed to a ‘two-hit’ stress model (MIA + loss of the social component of environmental enrichment (EE)), mice did not display anhedonia but required an increased number of PD and RL correction trials. These animals also had significant reductions of CamK2a mRNA in the prefrontal cortex. Appropriate functioning of synaptic plasticity, via mediators such as this protein kinase and others, are critical for behavioral flexibility. Although EE has been implicated in delaying the appearance of symptoms associated with certain brain disorders, these findings are in line with evidence that it also makes individuals more vulnerable to its loss. Overall, with the right ‘dose’, early life stress exposure can confer at least some functional advantages, which are lost when the number or magnitude of these exposures become too great.

## Introduction

Numerous epidemiological studies have identified an association between inflammatory insults during pregnancy and the prevalence of neuropsychiatric disorders, such as autism and schizophrenia, in offspring (Babulas et al., 2006; Estes and McAllister, 2016; Sørensen et al., 2008). However, at the population level, only a subset of children exposed to prenatal infections develop disease-relevant behavioral abnormalities (Mahic et al., 2017; Meyer, 2019). This suggests the etiology of these neurodevelopmental disorders may involve a synergism of prenatal immune challenge and other environmental risk factors (e.g., stress) in later life. Adding to this complexity, the effects of prenatal immune insults on child health outcomes also vary depending on the child’s biological sex (Gilman et al., 2016; Goldstein et al., 2014; Mac Giollabhui et al., 2019). Hence, it is essential to determine sources that contribute to the heterogeneous outcomes of MIA and to identify factors for building neurodevelopmental resilience and susceptibility to this early-life adversity.

In preclinical research, rodent maternal immune activation (MIA) models have been employed to mimic the effects of gestational infections and to investigate its underlying biological mechanisms (Estes and McAllister, 2016; Kentner et al., 2019a; Knuesel et al., 2014). Specifically, a mid-gestational injection of polyinosinic:polycytidylic acid (poly (I:C)), a viral mimetic toll-like receptor 3 agonist, can induce an extensive collection of innate immune responses (Mueller et al., 2019) and lead to a wide array of abnormalities in neurophysiology, behavior and cognitive abilities (see (Haddad et al., 2020) for review). Whereas the vast majority of previous studies have addressed the adverse effects of MIA, prior research has also revealed that MIA-treated offspring can exhibit improvements in cognitive functioning (Makinson et al., 2019; Nakamura et al., 2021), blunted responses to a second immune challenge in adolescence (Clark et al., 2019), and a higher level of resilience to the disruptive effects of isolation rearing on behavior and neurophysiology (Goh et al., 2020).

Recently, it has been shown that poly (I:C)-induced MIA at a critical window of parvalbumin interneuron development can lead to improvements in spatial working memory (Nakamura et al., 2021). These findings indicate that MIA may act like other mild early-life stressors to facilitate the development of protection against stressors in later life (Fujioka et al., 2001; Cannizzaro et al., 2006; Li et al., 2013). However, our understanding of the adaptations and potential priming effects of MIA on resiliency are relatively underexplored as the focus is more typically a deficit-based one. Notably, the ‘hidden talents’ approach has provided a framework for investigating how harsh and unpredictable childhood environments may promote changes in behaviors and cognition that are adaptive for future harsh environments in humans (see Ellis et al., 2020), explaining some of these seemingly paradoxical observations in animal studies. This is in line with the ‘stress acceleration hypothesis’ and the view that early life adversity accelerates the development of emotion based behaviors and neural circuits which may promote mental health resilience (see Callaghan & Tottenham, 2016). Together, the evidence surrounding these perspectives suggest that early life experiences influence a variety of brain circuits, affecting both emotional and cognitive functioning, which may have context-specific benefits.

The complexity of the living environment affects brain development at the functional, anatomical, and molecular level (Kempermann, 2019; Ohline and Abraham, 2019; Zhang et al., 2018). Increased complexity of the laboratory animal home cage, so-called environmental enrichment (EE), is characterized by exposure to environments with rich social, motor, cognitive and sensory stimulation. Although EE has been implicated in delaying the appearance of symptoms associated with certain brain disorders (Chourbaji et al., 2011; Herring et al., 2009; Van Dellen et al., 2000), it also makes individuals that are exposed to EE more vulnerable to its loss. Indeed, when the environment changes from being enriched to more impoverished (EE removal) behavioral alterations linked to depressive symptomology occur (Morano et al., 2019; Smith et al., 2017). Moreover, these behavioral changes are accompanied by dysregulation of the hypothalamus-pituitary-adrenal axis activity in female rats (Morano et al., 2019). EE removal may therefore be considered as a second hit factor which could potentially reinforce the disruptions induced by the first hit (e.g., MIA) in early-life and ultimately lead to onset of a full clinical syndrome.

In the present study, we aimed to examine how MIA interacts with the loss of the social component of life-long EE, influencing neuropathology and cognitive functioning of both male and female offspring. Social isolation has been implicated in mediating a range of behavioral deficits in adulthood including sensorimotor processing, social interaction abnormalities, heightened anxiety and cognitive dysfunction (Bakshi and Geyer, 1999; Fone and Porkess, 2008). By integrating this component, we can further explore mechanisms underlying the variation in the neurodevelopmental resilience and susceptibility to a ‘second-hit’ stressor in later life. Impairments in the cognitive functions were evaluated by leveraging a touchscreen-based operant task paradigm. Pair-wise discrimination (PD) and reversal learning (RL) tasks have been employed to assess impairments in motivation, memory, rule learning, and cognitive flexibility (Bryce and Howland, 2015; Bussey et al., 2012; Horner et al., 2013; Lins et al., 2018).

To examine the neuropathological alterations, we focused on the prefrontal cortex (PFC), an important brain region for cognitive control (Miller and Cohen, 2001). In this region, we measured the density of parvalbumin (PV+) cells and perineuronal nets (PNNs), which are involved in various functions including the regulation of signal transmission and circuit plasticity (Testa et al., 2019), and in supporting cognitive processes including working memory and attentional set shifting (Cho et al., 2015; Fuchs et al., 2007; Sohal et al., 2009). PV+ expression abnormalities and malformation of PNNs have been implicated in the etiology of neurodevelopmental disorders (Fung et al., 2010; Hashimoto et al., 2003; Testa et al., 2019). To further explore neurochemical alterations in the PFC, we analyzed mRNA expression of neural markers associated with different neurotransmitter pathways.

## 2. Materials and Methods

### 2.1. Animals

Male and female C57BL/6J mice arrived from the Jackson Laboratory (Bar Harbor, ME), and were housed at 20°C on a 12 h light/dark cycle (0700–1900 light) with ad libitum access to food and water. **Figure 1** details the timeline of experimental procedures which took place during the light phase. Female mice were housed as pairs in either larger environmental enrichment cages (EE; N40HT mouse cage, Ancare, Bellmore, NY), with access to toys, tubes, a Nylabone, Nestlets^®^ (Ancare, Bellmore, NY) and Bed-r’ Nest^®^ (ScottPharma Solutions, Marlborough MA), or standard sized cages (SD; N10HT mouse cage, Ancare, Bellmore, NY) with Nestlets^®^ and a Nylabone only. Males were paired in SD conditions until breeding. At that point, male animals were housed with two EE or SD females. Dams received either a 20 mg/kg intraperitoneal injection of polyinosine-polycytidylic acid (poly (I:C); tlrl-picw, lot number PIW-41-03, InvivoGen), or vehicle (sterile pyrogen-free 0.9% NaCl) on the morning of gestational day (G)12. Offspring were weaned into same-sex groups on P21 and maintained in their housing assignments (SD = 2-3 animals/cage; EE = 4-5 animals/cage) until P90. Additional methodological details, including the validation of poly (I:C), can be found in the reporting table from Kentner et al. (2019), provided as **Supplementary Table 1**. The MCPHS University Institutional Care and Use Committee approved all procedures described, which were carried out in compliance with the recommendations outlined by the Guide for the Care and Use of Laboratory Animals of the National Institutes of Health.

**Figure 1.**
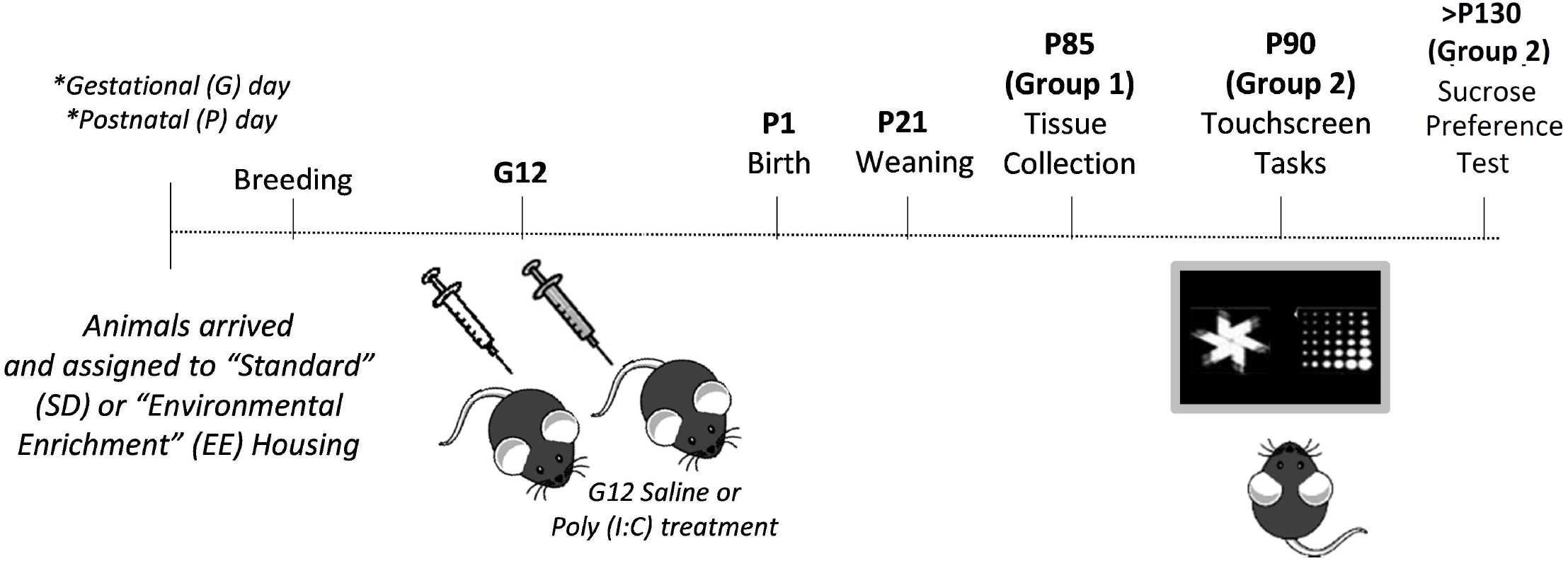
Outline of experimental procedures.

### 2.2. Touchscreen Learning

On P90, animals were individually housed in SD or EE conditions and food restricted (at 85-90% of their free-feeding weight which was maintained until the end of the study) to facilitate touchscreen responding for a milkshake reward (Strawberry Nesquik®). Prior to touchscreen training, animals were habituated to the liquid food reward, placed on a plastic dish, in their home cages (n = 8-13 litters represented per sex, MIA and housing group). Eight touchscreen sound-attenuated operant chambers (Campden Instruments Ltd, UK) were used to run the visual discrimination and reversal tasks. Details on the apparatus and dimensions have been reported previously (Horner et al., 2013). Training and testing were completed using the ABET II software.

Mice were first habituated to the chambers then trained to a) recognize the sound and light cues, b) interact physically with the touchscreen, and c) collect the liquid rewards following the manufacturer’s suggested protocol, and as previously outlined by Lins et al (2018). Chambers were thoroughly cleaned with Quat TB, and touchscreens with 10% ethanol alcohol (to maintain touch sensitivity), between each animal and session.

#### Visual discrimination

In this task, novel black and white images (the classic ‘fan’ and ‘marbles’) were simultaneously presented to mice (**Figure 1**). Left and right placement of the visual stimuli were presented in a pseudo randomized manner. For each animal, one of the two images was always correct (S+), regardless of its left vs right placement. When the animal responded to this S+ image, they received a continuously reinforced (100% probability of reward) delivery of ~7 ul of liquid milkshake. If the mouse responded to the incorrect (S-) image, no reward was produced and the animals had a 5-second ‘time out’ period, followed by a correction trial. Once the animal correctly responded to the correction trial (where the two images were presented in the same spatial configuration as the previous trial in which the animal had responded incorrectly), the next trial of the session commenced. The designation of the correct stimulus (fan vs marble) was counterbalanced across mice. Criterion was reached when the animal completed 30 trials in 1 hour (excluding correction trials) per daily session with >85% correct for two consecutive days. If an animal could not meet criterion after 30 days, they did not progress onto the next task (Radke et al., 2019; Kenton et al., 2020).

#### Reversal learning

Reversal learning was evaluated after successful completion of the visual discrimination task (n = 7-11 litters represented per sex, MIA and housing group). In reversal learning the previously correct image (S+) becomes the incorrect visual stimuli (S−) while the previously incorrect image (S−) becomes the correct choice (S+) that would result in a liquid reward delivery. Criterion was reached when animals reached 30 trials in 60 minutes with >85% correct for two consecutive days or were tested for 10 days (Prado Lab, 2019; Van den Broeck et al., 2019).

### 2.3. Sucrose Preference Test

To evaluate the effect of social enrichment loss on anhedonia, we utilized the 16-hr overnight sucrose preference test (Connors et al., 2014; Kentner et al., 2010). Here, male and female animals were given two bottles, each containing either a 1% sucrose solution or water. Sucrose preference was determined by calculating the ratio of sucrose intake (grams) to total fluid intake (grams) and converted into a percent score. The placement of the sucrose and water bottles was counterbalanced between trials to prevent side preferences. At no time were animals deprived of food or water.

### 2.4. Tissue collection and analysis

To circumvent the confound of touchscreen training modifying the brain, we collected tissue from littermates matched to our touchscreen animals on P85 (n = 8 litters represented per sex, MIA and housing group). This allows for a better assessment of the role of poly (I:C) in our data interpretation. A mixture of Ketamine/Xylazine (150 mg/kg, i.p/15 mg/kg, i.p) was used to anesthetize animals. Animals were perfused intracardially with a chilled phosphate buffer solution. Prefrontal cortex (PFC) and ventral hippocampus were dissected from the hemisphere, frozen on dry ice and stored at −75°C until processing. The other hemisphere was post-fixed in a 4% paraformaldehyde, phosphate buffer 0.1M solution (BM-698, Boston BioProducts) overnight at 4°C. Tissue was submerged in ice cold 10% sucrose in PBS (with 0.1% sodium azide) and incubated at 4°C overnight. The next day, solution was replaced with 30% sucrose in PBS for 3 days. Tissue was then flash frozen with 2-methylbutane (O3551-4, Fisher Scientific) and stored at −75°C until sectioning.

### 2.5. RT-PCR

Using the RNeasy Lipid Tissue Mini Kit (Qiagen, 74804), total RNA was extracted from frozen tissue and resuspended in RNase-free water. A NanoDrop 2000 spectrophotometer (ThermoFisher Scientific) was used to quantify isolated RNA. Total RNA was reverse transcribed to cDNA with the Transcriptor High Fidelity cDNA Synthesis Kit (#5081963001, Millipore Sigma) using the manufacturers protocol, and the final cDNA solution was stored at −20°C until analysis. Quantitative real-time PCR with Taqman™ Fast Advanced Master Mix (#4444963, Applied Biosystems) was used to measure the mRNA expression (See **Table 1** for list of genes expression assays). Reactions were analyzed in triplicate using optical 96-well plates (Applied Biosystems StepOnePlus™ Real-Time PCR System) and relative gene expression levels were evaluated using the 2 ^−^ΔΔ^CT^ method with 18S (Hs99999901_s1). Gene expression was normalized in relation to 18S and presented as mean expression relative to same sex SD-saline treated controls. As shown previously, 18S did not differ across groups (Zhao et al., 2021).

**Table 1.**
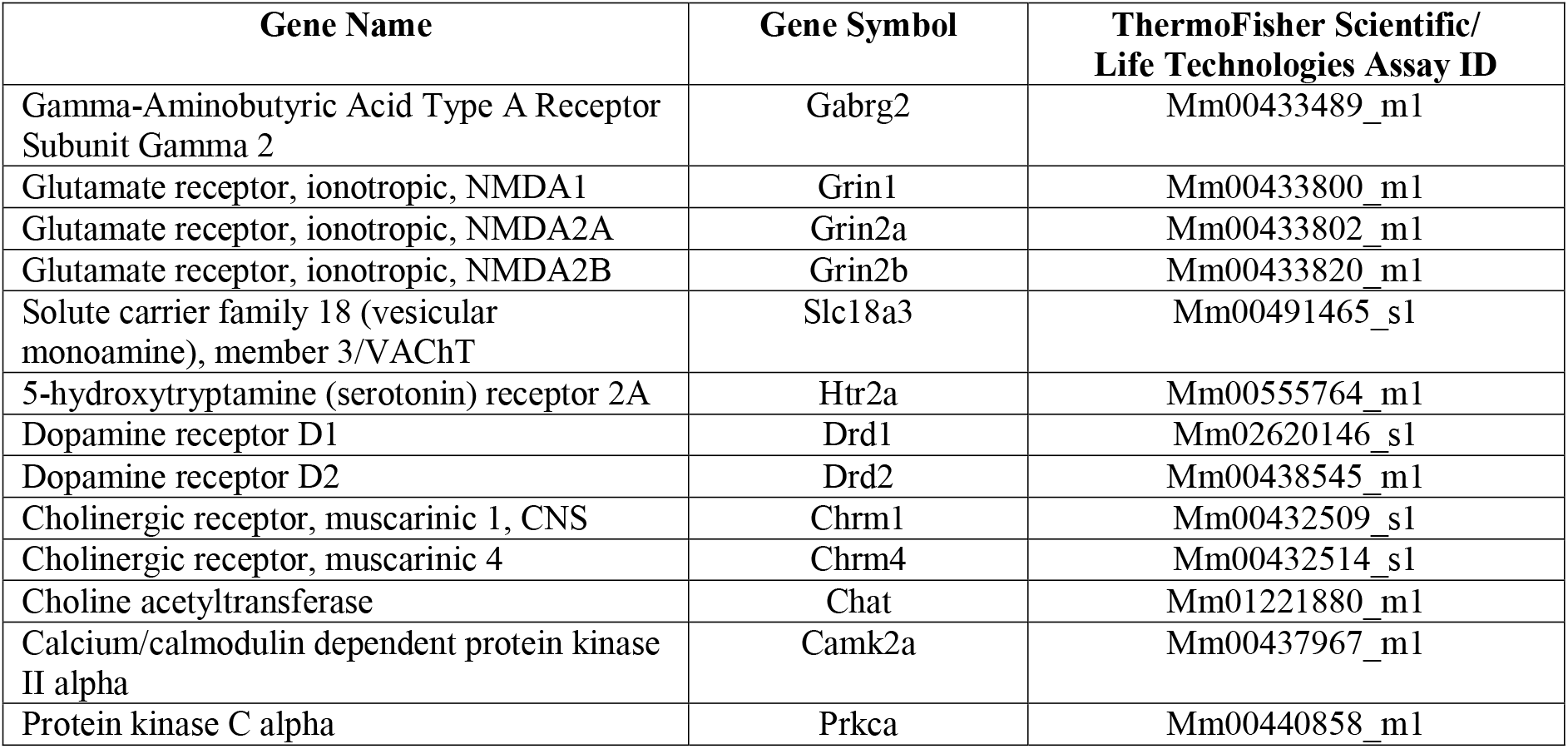
Gene Expression Assays for TaqMan™ qPCR

### 2.6. Immunohistochemistry

Coronal sections (40 um thick) were obtained on a cryostat (Leica CM1860) as a 1:4 series and stored at −20°C in cryoprotectant until immunocytochemistry. Free-floating sections were washed in PBS to remove the cryoprotectant. Prefrontal cortex sections were selected from 2.09 to 1.93 mm caudal to Bregma based on the Paxinos & Franklin, 2019 brain atlas. For tissue processing, sections were mounted onto charged slides and dried overnight. Slides were then rinsed with PBS and allowed to incubate in primary antibodies for wisteria floribunda agglutinin (PNNs; 1:1,000; Sigma-Aldrich L1516) and parvalbumin (PV+; 1:500; RnD cat# AF5058) for 1 hour at room temperature, followed by 48 hours at 4 degrees C. Slides were then washed with PBS, treated with secondary antibodies Alexafluor 594 (1: 600; cat# A-11016) and Alexafluor 488 (1: 600; cat#S112233) in PBS-T, washed again with PBS, and immediately cover slipped with Vectashield antifade mounting medium (Vector Laboratories H-1500-10). Sections were imaged at 20x magnification and stitched on the Keyence BZ-X800 all-in-one fluorescence microscope for counting of the prelimbic cortex (PL), infralimbic cortex (IL), medial orbital cortex (MO) and ventral orbital cortex (VO). There were 1 to 3 sections taken for each brain region of interest. For example, MO only appeared on one plate in the brain atlas, so there was one section per animal counted for this region. VO appeared on two plates in the brain atlas, so there were two sections per animal counted for this region. The number of both PV+ and PNNs were counted manually in ImageJ by two researchers blind to the experimental groups (inter-rater reliability >0.90). A PNN was identified by being a fully formed “halo,” and any partial PNN or background staining was not counted.

### 2.7. Statistical analyses

The statistical software package Statistical Software for the Social Sciences (SPSS) version 26.0 (IBM, Armonk, NY) was used for all analyses, except for the survival curves which were evaluated with the Log-rank (Mantel Cox test) using GraphPad Prism (Version 9.0). Following the overall omnibus test evaluating all eight groups, individual Log-rank tests were conducted as post hocs, alongside a Bonferonni-Holm alpha correction to protect against multiple comparisons (Giacalone et al., 2018).

Two or three-way ANOVAs were used as appropriate. In cases of significantly skewed data (Shapiro-Wilks), Kruskal-Wallis tests (non-parametric equivalent to ANOVA, reported as Χ^*2*^) were applied. Body weight was used as a covariate for sucrose preference data. LSD post hocs were applied, except where there were fewer than three levels in which case pairwise t-tests and Levene’s were utilized. Multiple testing procedures for the gene expression data were analyzed using the False Discovery Rate (FDR). Partial eta-squared (*n_p_*^2^) is also reported as an index of effect size (the range of values being 0.02 = small effect, 0.13 = moderate effect, 0.26 = large effect; Miles and Shevlin, 2001).

## 3. Results

### 3.1. Behavioral tests

#### 3.1.1 Touchscreen

The overall Log-rank (Mantel-Cox) test was significant across all groups, suggesting a difference in session-to-criterion survival curves (X^2^ (7) = 19.82, p = 0.006; **Figure 2A**). Post hoc tests failed to identify differences across MIA, housing, or sex using adjusted alpha values that accounted for the number of comparisons made (p>0.01). However, ANOVA revealed a main effect of sex on number of sessions required to reach criterion in the discrimination task (F(1, 67) = 12.086, p = 0.001, *np*^2^ = 0.153; male: 9.94±0.82 vs female: 15.9±1.38).

**Figure 2.**
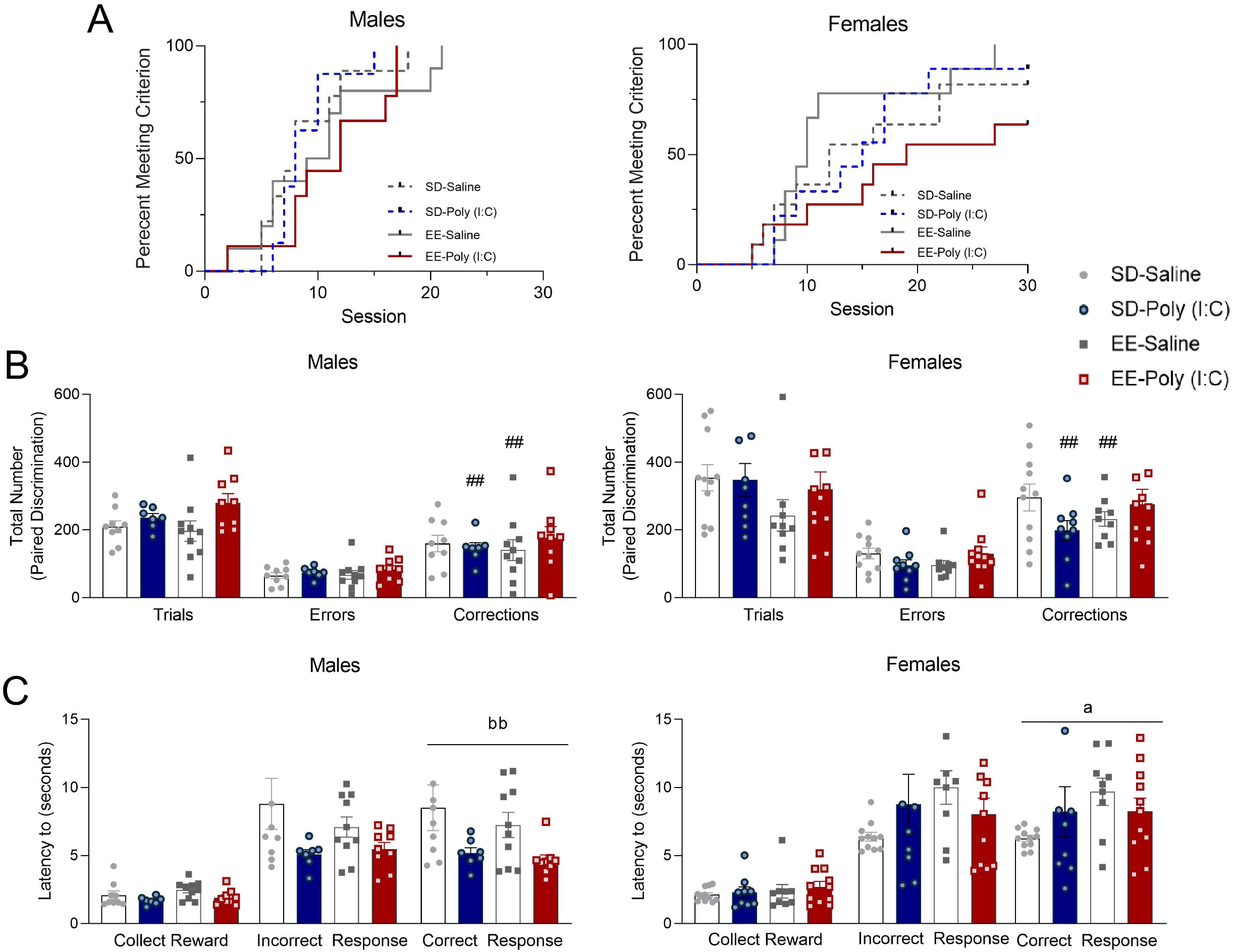
Paired visual discrimination learning in male (left) and female (right) MIA offspring. (A) Survival analysis of sessions to discrimination criterion (n = 8-13 litters represented per sex, MIA and housing group). (B) Total number of trials, errors, and correction trials completed by animals that met criterion during all discrimination sessions. (C) Latency (seconds) to respond for a correct choice, incorrect choice, and reward during all discrimination sessions. Data expressed as mean ± SEM, n=7-11 litters represented per sex, MIA, and housing group. ##p < 0.01, versus EE-poly (I:C); ^a^p < 0.05, main effect of housing; ^bb^p < 0.01, main effect of MIA.

The total number of discrimination trials and errors did not differ across groups (p>0.05; **Figure 2B**). A MIA by housing interaction (F(1,67) = 4.028, p = 0.049, *np*^2^ = 0.057) revealed that EE-poly (I:C) mice required more correction trials compared to their EE-saline (t(37) = −3.416, p = 0.002) and SD-poly IC counterparts (t(34) = −2.788, p = 0.009). This suggests that the combination of multiple early life stressors (MIA + companion loss) compounded impairments on this measure (**Figure 2B**). Latency to collect rewards and make incorrect responses was not different as a function of sex, MIA or housing (p>0.05; **Figure 2C**). However, there were significant sex differences in the average latency to make a correct response in the visual discrimination task (*X*^2^ (1) = 5.311, p = 0.021). Specifically, male treated MIA (*X*^2^ (1) = 6.501, p = 0.011; poly (I:C): 4.89 ± 2.78 vs SD: 7.84 ± 0.92) mice were faster, and female EE ((*X*^2^ (1) = 4.116, p = 0.042; SD: 7.14 ± 0.84 vs EE: 8.89 ± 0.704) mice were slower, to respond when making a correct choice (**Figure 2C)**. In alignment with other research suggesting that females thrive better in social environments, this suggests that female EE mice were more sensitive to the loss of companionship in their homepage.

For reversal learning, there were no group differences in session-to-criterion survival curves (Long-rank test, X^2^ (7) = 10.26, p = 0.17 (N.S); **Figure 3A**) or in the number of sessions to reach reversal criterion (p>0.05). The total number of trials completed were lower for female poly (I:C) animals relative to their saline counterparts (sex x MIA: F(1, 56) = 14.064, p = 0.001, *np*^2^ = 0.201; female poly (I:C) vs female saline (p = 0.002); **Figure 3B**). Additionally, female poly (I:C) mice made fewer errors compared to female saline animals (sex x MIA: F(1, 56) = 11.24, p = 0.001, *np*^2^ = 0.167; female poly (I:C) vs female saline (p = 0.005); **Figure 3B**) suggesting that early life challenges may not only lead to impairments, but may be associated with some performance benefits. A significant sex x housing x MIA interaction (F(1, 57) = 7.048, p = 0.010, *np*^2^ = 0.110) showed that male (p = 0.027) and female (p = 0.0001) SD-poly (I:C) animals required fewer correction trials than same-sex EE-poly (I:C) mice. This is in line with evidence that multiple hits can result in cognitive deficits that are not apparent with only one stressor exposure (Bilbo et al., 2005). Moreover, female SD-poly (I:C) mice had fewer correction trials than SD-saline animals (p = 0.0001; **Figure 3B**), again highlighting the potential for early life adversity to confer functional advantages. There were no differences in average latency to collect rewards or to make incorrect and correct responses on the reversal trials (p >0.05; **Figure 3C**).

**Figure 3.**
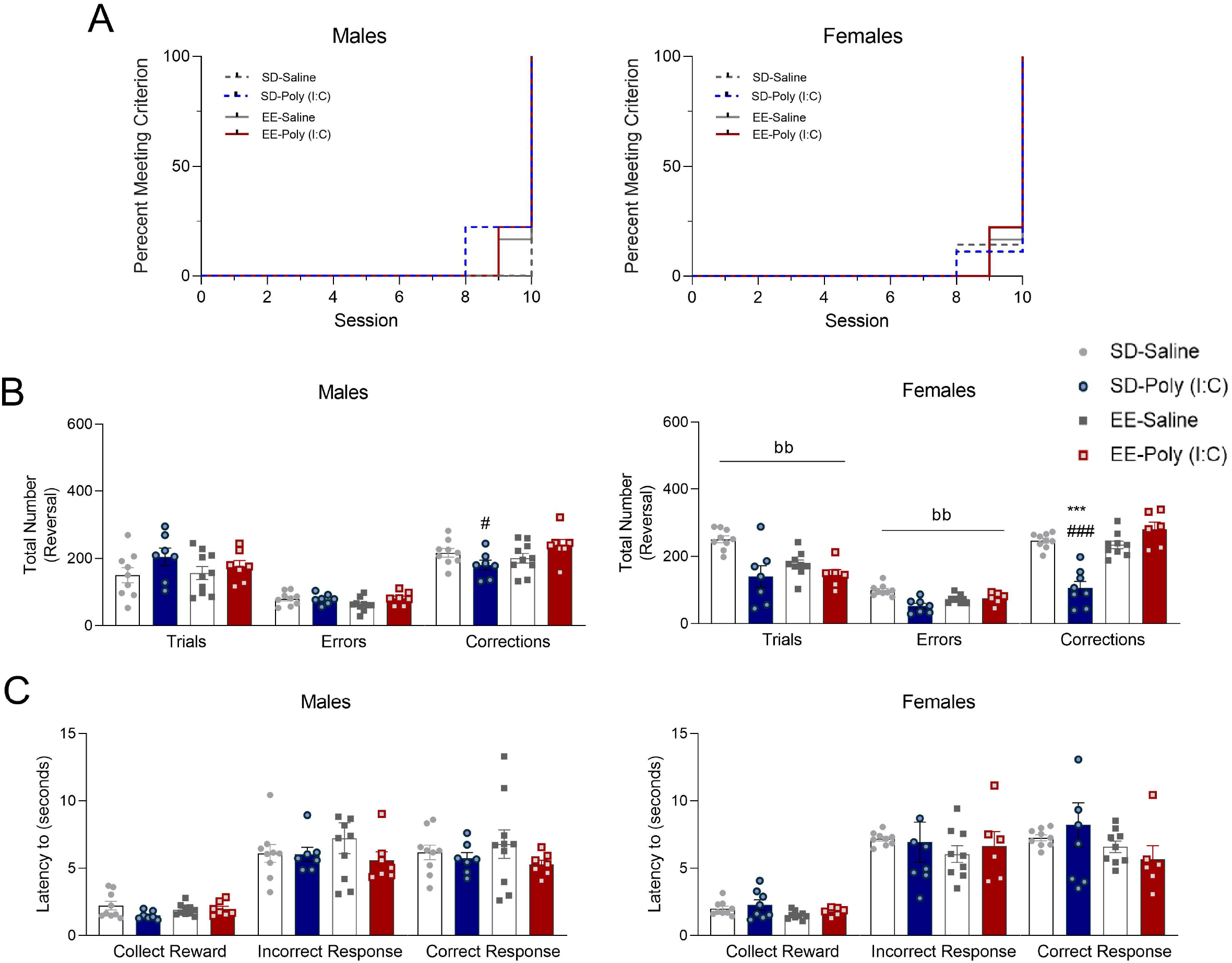
Reversal learning in male (left) and female (right) MIA offspring. (A) Survival analysis of sessions to reversal criterion. (B) Total number of trials, errors, and correction trials completed during all reversal sessions. (C) Latency (seconds) to respond for a correct choice, incorrect choice, and reward during all reversal sessions. Data expressed as mean ± SEM, n=7-11 litters represented per sex, MIA, and housing group. ***p < 0.001, versus SD-saline; #p < 0.05, ###p < 0.001, versus EE-poly (I:C); ^bb^p < 0.01, main effect of MIA.

#### 3.1.2. Sucrose Preference Tests

To determine whether EE-poly (I:C) animals had motivational, rather than cognitive, impairments associated with the multiple stressor experience (MIA + social companion loss) we conducted a 16-hour sucrose preference test. We did not observe anhedonia using this measure (p>0.05; **Supplementary Figure 1**). Moreover, animals were not different in their latency to collect the milkshake rewards (**Figure 2C; Figure 3C**). Therefore, we did not find motivational deficits in these animals that would interfere with their touchscreen responding.

### 3.2. Taq Man qPCR

#### 3.2.1. GABA and Glutamatergic Receptor Expression

We observed that levels of Gabrg2 in the PFC were elevated in poly (IC) treated (F(1, 28) = 5.007, p = 0.033, *np*^2^ = 0.152; poly (IC): 1.10±1.09 vs saline: 0.85±0.07) and standard housed mice (males: F(1, 28) = 10.768, p = 0.003, *np*^2^ = 0.278; SD: 1.16±0.09 vs EE: 0.79±0.09; females: F(1, 28) = 9.606, p = 0.004, *np*^2^ = 0.255; SD: 1.09±0.05 vs EE: 0.82±0.07; **Supplementary Table 2**). Grin1 expression was increased in female SD-poly (IC) animals compared to SD-saline (p = 0.004) and EE-poly (IC) mice (p = 0.003; housing x MIA interaction: F(1, 28) = 13.250, p = 0.001, *np*^2^ = 0.321; **Figure 4A**). There were no housing or MIA effects on Grin2a (p>0.05; **Supplementary Table 2**), however PFC levels of Grin2b were elevated in male poly (IC) mice (F(1, 28) = 21.745, p = 0.0001, *np*^2^ = 0.437; poly (IC): 1.28±0.52 vs saline: 0.95±0.04; **Figure 4B**). Additional statistical analyses of ventral hippocampal levels of each of these genes can be found in **Supplementary Table 2**.

**Figure 4.**
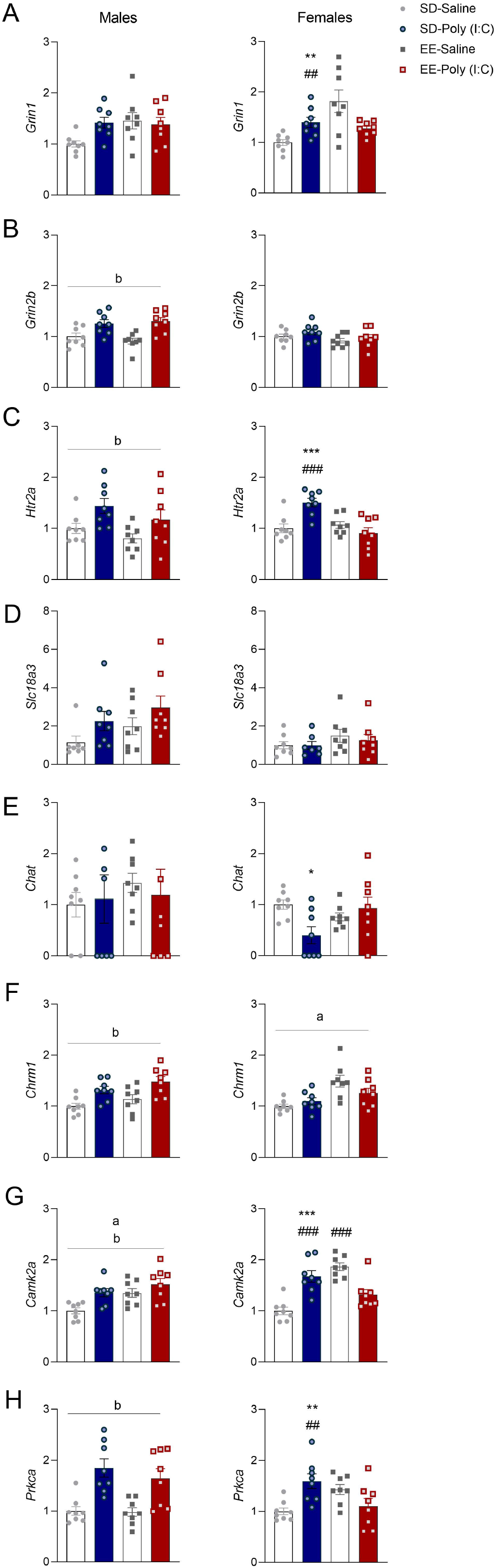
Prefrontal cortex gene expression in male (left) and female (right) MIA offspring on postnatal day 85. Levels of (A) Grin1, (B) Grin2b, (C) Htr2a, (D) Slc18a3 (E) Chat, (F) Chrm1, (G) Chrm4, and (H) Prkca mRNA. Gene expression data are expressed as mean ± SEM, n=8 litters represented per sex, MIA, and housing group. *p < 0.05, **p < 0.01, versus SD-saline; ##p < 0.01, versus EE-poly (I:C).

#### 3.2.2. Markers of Monoamine and Cholinergic Functioning

A variety of receptor mechanisms involved in monoaminergic and cholinergic signaling are critical for PFC mediated cognitive functioning (Hupalo et al., 2019; Waltz, 2017). While male SD mice had higher Drd1 expression levels compared to EE animals in the PFC (p = 0.017; main effect of housing: F(1, 28) = 6.458, p = 0.017, *np*^2^ = 0.187; SD: 1.109±0.07 vs EE: 0.85±0.07; **Supplementary Table 2**), there were no significant group effects of Drd2 (p>0.05; this and ventral hippocampal dopamine receptor genes are provided in **Supplementary Table 2**). In contrast, Htr2a levels in PFC were elevated in poly (I:C) males compared to saline (p = 0.007; main effect of MIA: F(1, 28) = 8.484, p = 0.007, *np*^2^ = 0.233; poly (I:C): 1.304±0.12 vs saline: 0.90±0.07; **Figure 4C**). SD-poly (I:C) was also associated with increased Htr2a in the PFC of female mice compared to SD-saline (p = 0.001) and EE-poly (I:C) animals (p = 0.001; MIA x housing interaction: F(1, 28) = 14.615, p = 0.001, *np*^2^=0.343; **Figure 4C**).

There were no group effects (p>0.05) for the vesicular acetylcholine transporter Slc18a3 (**Figure 4D**) or choline acetyltransferase mRNA (Chat; **Figure 4E**) in the PFC.

The muscarinic receptors Chrm1 and Chrm4 have both been implicated in the pathogenesis of schizophrenia (Gibbons & Dean, 2016). Chrm4 was significantly reduced in EE-saline vs EE-poly (I:C) male, but not female, mice in the PFC (p = 0.0001; MIA x housing interaction: F(1, 28) = 4.909, p = 0.035, *np*^2^= 0.149; **Supplementary Table 2**). While Chrm1 was elevated in female EE compared to SD mice (p = 0.001; main effect housing: F(1, 28) = 14.893, p = 0.001, *np*^2^ = 0.347; EE:1.37±0.08 vs SD: 1.05±0.03; **Figure 4F**), in males there was a significant main effect of MIA (F(1, 28) = 16.688, p = 0.0001, *np*^2^ = 0.373). Here, male poly (I:C) had elevated PFC levels of this muscarinic receptor compared to saline (p =0.0001; poly (I:C): 1.40±0.06 vs saline: 1.07±0.05; **Figure 4F**).

#### 3.2.3. Synaptic Plasticity Markers

Appropriate functioning of synaptic plasticity in the prefrontal cortex, via mediators such as CaMKII and other protein kinases, is critical for behavioral flexibility (**Ma et al., 2015; Natividad et al., 2015)**. In males, both Camk2a (**Figure 4G**) and Prkca (**Figure 4H**) expression levels were elevated in poly (I:C) compared to saline treated mice (main effect of MIA; CamK2a: F(1, 28) = 9.320, p = 0.005, *np*^2^ = 0.250; poly (I:C): 1.44±0.07 vs saline: 1.17±0.07; Prkca: F(1, 28) = 27.233, p = 0.0001, *np*^2^ = 0.493; poly (I:C): 1.75±0.13vs saline: 0.99±0.06). SD-poly (I:C) females had heightened expression of Camk2a (MIA x housing interaction: F(1, 28) = 43.633, p = 0.001, *np*^2^=0.609) and Prkca (F(1, 28) = 14.416, p = 0.005, *np*^2^ = 0.340) compared to SD-saline (Camk2a: p= 0.001; Pkrca: p = p = 0.002) and EE-poly (I:C) animals (Camk2a: p = 0.0001; Prkca: p = 0.003). EE-saline animals also had increased levels of Camk2a (p = 0.001) compared to EE-poly (I:C) animals, suggesting that multiple stressor hits may inhibit some of the advantages conferred by stressful experiences.

#### 3.2.4. Immunohistochemistry of PV+ and PNNs

Using a poly (I:C) rat model of MIA, previous investigators (Paylor et al., 2016) have found selective reduction of PNNs and the percentage of PV+ cells surrounded by PNNs in the medial prefrontal cortex region. We have extended these observations from the PL and IL to also evaluate MO and VO in mice. **Figure 5A** provides the representative plates with our PFC regions of interest outlined while **Figure 5B** shows representative PV+, PNN, and PV+/PNN colocalization of cells. With respect to the PL, EE male and female animals had higher expression of PV+ cells (SD: 25.36 ± 2.55; EE: 33.70 ± 2.62; F(1, 42) = 5.225, p = 0.027, *np*^2^=0.111: **Figure 5C**). There was a MIA effect in this region in that the density of PNNs was significantly reduced in both male and female MIA mice (Saline: 10.40 ± 0.99; poly (I:C): 7.40 ± 0.62; F(1, 42) = 6.330, p = 0.016, *np*^2^=0.131: **Figure 5C**). While the percent of colocalized PV+ cells ensheathed in PNNs did not vary for males in the PL (p>0.05), there was a significant MIA by housing interaction for female mice (F(1, 42) = 7.193, p = 0.010, *np*^2^=0.146; **Figure 5C**). Specifically, EE-saline female mice had a higher percentage of colocalization than SD-saline (0.045) and EE-poly ((I:C); p = 0.003) animals in this brain region.

**Figure 5.**
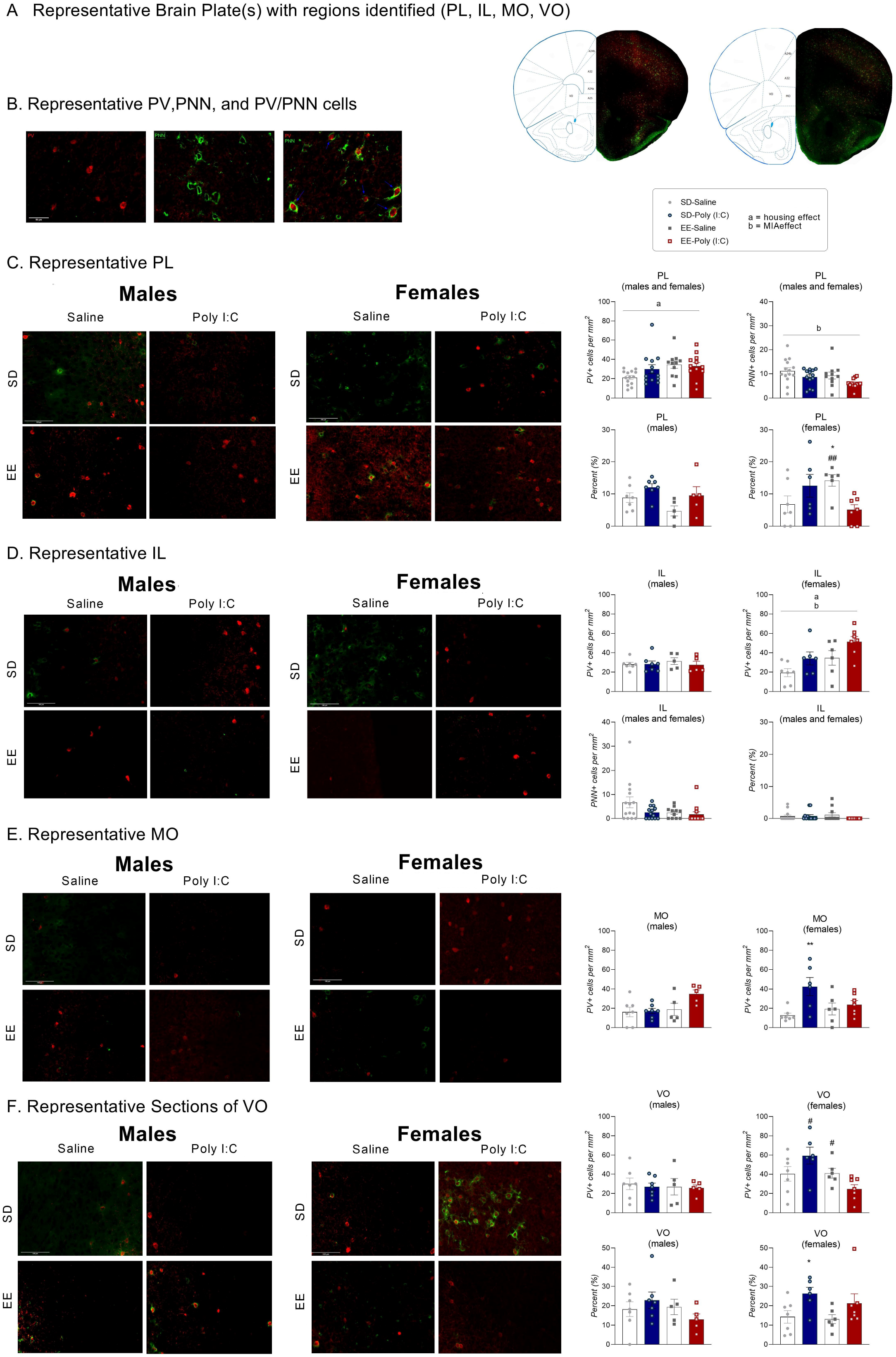
Expression of parvalbumin (PV+) and perineuronal net (PNN) density for male and female MIA offspring on postnatal day 85. (A) Representative brain plates for brain regions of interest such as prelimbic (PL), infralimbic (PL), medial orbital (MO) and the ventral medial (VO) cortices. (B) Representative depictions of PV+, PNN, and PV/PNN colocalization. Representative sections and associated data for (C) prelimbic, (D) infralimbic, (E) medial orbital and (F) ventral orbital cortex. Data are expressed as mean ± SEM; n=5-7 litters represented per sex, MIA, and housing group. Where there were no sex differences, male and female data were collapsed for visualization purposes. *p < 0.05, **p < 0.01, versus SD-saline; #p < 0.05, ##p < 0.01, versus EE-poly (I:C). ^a^p < 0.05, main effect of housing; ^b^p < 0.05, main effect of MIA.

In terms of the IL, there were significant interactions between sex and MIA (F(1, 42) = 6.525, p = 0.014, *np*^2^=0.134) and sex and housing (F(1, 42) = 6.429, p = 0.013, *np*^2^ = 0.133; **Figure 5D**). Female MIA mice had higher expression of PV+ positive cells in the IL compared to saline treated (saline: 26.465 ± 4.543; poly (I:C): 43.494 ± 4.774; t(24) = 2.584, p = 0.016). Enriched housed females were found to demonstrate the same pattern of PV+ expression compared to SD (SD: 26.193 ± 4.233; EE: 43.766 ± 4.974; t(24) = −2.691, p = 0.013). No significant interactions or main effects were observed for either density of PNNs or percent of colocalization of PV+ and PNNs in the IL (**Figure 5D**; p>0.05).

There was a significant sex by MIA by housing interaction (F (1, 42) = 7.207, p = 0.010, *np*^2^ = 0.146) for PV+ expression in the MO where male SD-poly (I:C) treated rats had lower expression compared to EE-poly (I:C) mice (t(10) = −3.765, p = 0.004; **Figure 5E**). Female SD-poly (I:C) mice had elevated expression of PV+ cells compared to SD-saline controls (t(11) = 3.339, p = 0.007; **Figure 5E**). In the MO, density of PNNs was higher in male (6.48 ± 1.09) compared to females (3.67 ± 0.74; F(1, 42) = 4.411, p = 0.042, *np*^2^ = 0.095) and in SD (5.35 ± 2.03) versus EE housed mice (2.29 ±1.59; *X*^2^(1)=6.166, p = 0.013; **Figure 5E**).

The VO region was also associated with a significant sex by MIA by housing interaction for PV+ expression (F(1, 42) = 4.360, p = 0.043, *np*^2^ =0.094). While follow-up analyses did not find any differences in males (p>0.05; **Figure 5F**), female EE-poly (I:C) mice had lower PV+ expression compared to SD-poly ((I:C); t(11) = −3.674, p = 0.004) and EE-saline animals (t(11) = −2.426, p = 0.034; **Figure 5F**). Post hoc tests following a significant sex by MIA by housing interaction for PNNs (F(1, 42) = 5.095, p = 0.029, *np*^2^ =0.108) did not detect group differences (p>0.05). However, while a sex and MIA interaction (F(1, 42) = 4.122, p = 0.049, *np*^2^ =0.089) for the percentage of PV+ cells ensheathed in PNNs revealed no differences among male mice (p>0.05; **Figure 5F**), female SD-poly (I:C) mice had a significantly higher percentage of colocalization compared to SD-saline animals of the same sex (t(11) = 2.617, p = 0.024; **Figure 5F**). Overall, there were heterogeneous changes in expression of PV+ and PNNs throughout the PFC region, as a function of both MIA and housing.

## Discussion

In the current study, we demonstrate that poly (I:C)-induced MIA imposes sex-specific enhancements on a variety of learning motifs, which stands in line with recent findings on enhanced spatial working memory in prenatally treated poly (I:C) mice (Nakamura et al., 2021) and improved cognitive performance in the 5-choice serial reaction time task in LPS-mice treated prenatally (Makinson et al., 2019). At first glance, these findings stand in contrast to previous studies reporting disrupted performance in the working memory, executive function, and cognitive flexibility of these animals (Amodeo et al., 2019; Lins et al., 2018; Meehan et al., 2017; Meyer et al., 2010; Murray et al., 2017; Wallace et al., 2014; Gogos et al., 2020). In addition to the differences in behavioral tasks used across these studies, it should be noted that several factors such as the immune stimulus intensity and the timing of challenge may also contribute to the discrepancies between the findings. For example, compared to earlier poly (I:C) exposure, MIA applied at later gestational stages (e.g. GD 14.5 and 17.5) is more influential on the display of schizophrenia-related behaviors (Meehan et al., 2017).

Our results suggest that MIA in mid-gestation may have an enhancing effect on certain aspects of cognitive functions. This echoes the notion of the ‘hidden talents’ approach (Ellis et al., 2020) which suggests that improved cognitive abilities may be an adaptive mechanism for individuals who grew up in harsh and unpredictable environments. The improved abilities on some measures are hypothesized to support success within predicted future environments of the same difficult nature. This ‘hidden talents’ framework could help explain some of these seemingly paradoxical observations that are arising in the MIA model. Alongside the commonly used ‘deficit-based’ approach, it could be a complementary perspective for considering the differential effects of many types of early experiences on later-life functioning (Ellis et al., 2020).

Sex differences in cognitive abilities have been well studied in both humans and animal models (review see (Hyde, 2016; Jonasson, 2005) and we found in the current study that male mice required fewer sessions than females to reach criterion in the PD task. Generally, touchscreen-based PD and RL tasks are used to assess impairments in motivation, memory, rule learning, and cognitive flexibility (Bryce and Howland, 2015; Bussey et al., 2012; Horner et al., 2013; Lins et al., 2018) and the sex differences uncovered here may reflect a small male advantages in some aspects of these abilities (Berger-Sweeney et al., 1995; Jonasson, 2005). However, it should be mentioned that in addition to showing a reduced latency to make a correct response, male MIA mice also seemed to show a reduced latency to make an incorrect response. Although this latter effect was non-significant, it is suggestive of a possible impulsivity phenotype in these males. Moreover, our female SD-poly (I:C) mice required fewer correction trials than SD-saline controls to complete RL task. This suggests that the enhancing effect of MIA exposure may benefit females more than males. This is consistent to previous work showing that in the 5-choice serial reaction time task, LPS-treated females learned the task more quickly and displayed a higher level of motivation to earn a reinforcer (decreased time required to reach criterion; Makinson et al., 2019). It is also noted that the sex dependent effects of MIA on cognitive performance may also be species and strain dependent (Gogos et al., 2020; Lins et al., 2018; Lins et al., 2019; Nakamura et al., 2021).

Long-term EE exposure followed by the loss of social enrichment may dampen the enhancing effects of MIA. This is supported by our findings that male and female SD-poly (I:C) animals required fewer correction trials than same-sex EE-poly (I:C) mice. In the current study, the cognitive tasks were conducted following social isolation, which has been implicated in mediating a range of behavioral deficits in adulthood including sensorimotor processing, social interaction abnormalities, heightened anxiety and cognitive dysfunction (Bakshi and Geyer, 1999; Fone and Porkess, 2008). The social isolation was a component of the standard protocol for touchscreen training, to facilitate food restriction to increase an animal’s responding for food rewards (Horner et al., 2013). Although both SD and EE animals were subjected to social isolation, the experience appeared to impose negative impacts on the cognitive performance of EE-poly (I:C) mice. This is consistent with the notion that loss of enrichment may elicit unique changes in behavioral and physiological phenotypes indicative of stress or despair (Morano et al., 2019). Specifically, single housing following prolonged EE exposure was associated with increased weight gain, elevated helplessness and passive coping behaviors, and blunted hypothalamic-pituitary-adrenocortical activity (Morano et al., 2019; Smith et al., 2017). Social isolation itself causes structural changes (e.g., impaired myelination and decreased density of dendritic spines) in the PFC (Lie et al., 2012; Silva-Gomez et al., 2003). In contrast, isolation rearing following MIA in SD-housed animals was protective against negative impacts on behavior and neurophysiology (Goh et al., 2020).

At the neurophysiological level, the mechanisms underlying the enhancing effects of MIA remain to be elucidated. Our current results suggest that protein kinase C (PKC) signaling in the PFC may be a potential pathway for mediating these effects. PKC isoforms are critically involved in many types of learning and memory (Alkon et al., 2007; Nelson et al., 2008; Sun and Alkon, 2010). Moreover, PKC can be activated by various synaptic inputs and intracellular signals that are involved in modulating cognitive functions, including glutamatergic (Hasham et al., 1997), cholinergic (Chen et al., 2005; Xiong et al., 2019) and serotonergic inputs (Carr et al., 2002; Carr et al., 2003). Therefore, the enhancing effects of MIA on cognition may be mediated through these inputs on the PKC signaling pathway. This is supported by our data showing upregulated *Prcka*, *Grin2b*, *Chrm1* and *Htr2a* in the poly (I:C)-treated male mice.

Changes in the balance between inhibition and excitation within the PFC, mediated through GABA and glutamate signaling, can impair behavioral flexibility (Bissonette & Powell, 2012; Enomoto et al., 2011). Calcium/calmodulin-dependent protein kinase type II subunit alpha (CamK2a) interacts with NMDA subunits and together they have a critical role in plasticity and learning (Ma et al., 2015; Takahashi et al., 2009). Given the parallel increase of these markers in our poly (I:C) animals (Group 1), we originally predicted that this upregulation was a strategy enacted by the brain in an attempt to compensate for the early life inflammatory challenge, but that we would still observe functional deficits in touchscreen performance (Group 2). Instead, the MIA animals surprised us by demonstrating sex-specific improvements in their PD and RL tasks. Notably, the upregulation of Grin2b expression in male PFC could be an indication of region-specific compensatory plasticity following MIA, since decreased levels of this receptor are reported in other brain regions and with other immunogens (Connors et al., 2014). The specific upregulation in Grin1 may also similarly facilitated improved learning in our female SD-poly (I:C) animals.

Given that poly (I:C) induced MIA in rodents is considered a model for schizophrenia (Meyer, 2019), and the role of muscarinic receptors in cognitive functioning, we were interested in their expression in this model. Chrm1 is thought to be important for synaptic plasticity (Gibbons & Dean, 2016) and treatment with selective agonists for this receptor have resulted in improvements in performance in spatial memory tasks (Vanover., et al., 2008), reversal learning (Shirey et al., 2009), and novel object recognition (Bradley et al., 2010). Our data of elevated PFC Chrm1 expression, alongside improved behavioral flexibility in male SD-poly (I:C) mice, are in line with these findings.

In contrast, the contribution of the 5HT2a receptor to cognition is complicated by conflicting reports of its antagonism both enhancing (Amodeo et al., 2014, Baker et al., 2011) and inhibiting behavioral flexibility (Boulougouris et al., 2008), and of its activation impairing probabilistic reversal learning (Amodeo et al., 2020). This suggests that the MIA-induced upregulation of PFC Ht2a in the current study should be associated with a cognitive impairment. Instead, we observed MIA mice to have a general improved performance in PD and RL touchscreen tasks. A critical difference in our work could be that the elevated receptor level in our female SD-poly (I:C) mice is likely reflective of a chronic, rather than acute, change. It is also possible that pharmacological 5Ht2a receptor activation may improve behavioral flexibility at different doses and more targeted central administration routes.

Both parvalbumin (PV+) and perineuronal nets (PNNs) are involved in the regulation of developmental critical periods in the cortex (Huang et al., 1999; Hensch, 2005; Takesian & Hensch, 2013). The fast-spiking inhibitory PV+ interneurons are frequently surrounded by PNNs, which provide metabolic support from the demand of these cells (Reinert et al., 2003; Härtig et al., 1999). Our results suggest that the altered expression of PV+/PNNs in the prefrontal cortex following MIA may contribute to the improved cognitive performance in female mice. Several previous studies have demonstrated that prefrontal PV+ interneurons support cognitive flexibility tasks through modulating gamma oscillations (Cardin et al., 2009; Cho et al., 2015; Sohal et al., 2009). In our current study, when compared to the saline controls, poly (I:C) females expressed higher levels of PV+ cells in the infralimbic cortex. SD-poly (I:C) females also expressed more PV+ cells in the medial orbital frontal cortex and more PV+ cells ensheathed with PNNs in the ventral orbital frontal cortex. By enwrapping the PV+ interneurons, PNNs can modulate their functions and protect against oxidative stress (Berretta et al., 2015; Favuzzi et al., 2017). Also, both infralimbic and orbital frontal cortex have been implicated in reversal learning (Avigan et al., 2020; Boulougouris et al., 2007; Chudasama and Robbins, 2003; Hervig et al., 2020; Leite-Almeida et al., 2014; Mukherjee and Caroni, 2018). We therefore speculate that the enhanced expression of PV+ cells in these regions may contribute to the improved reversal learning in the female MIA mice.

Besides gating PV+ interneuron function, PNNs are also involved in regulating experience-dependent circuit plasticity through enhanced synapse stabilization and by restricting synaptic rearrangements beyond the critical period for plasticity (Pizzorusso et al., 2002). Moreover, deficits of PNNs in the PFC have been observed in individuals with schizophrenia (Enwright et al., 2016; Mauney et al., 2013). In the current study, we found that MIA also led to reductions of PNNs in the prelimbic cortex of poly (I:C) offspring, which is consistent to a previous study using a rat MIA model (Paylor et al., 2016). However, future studies are needed to examine how these changes may lead to behavioral abnormalities, or even improved abilities on some measures. Overall, there were heterogeneous changes in the expression of PV+ and PNNs throughout the PFC regions, as a function of both MIA and housing. This is in line with the perspective that some neural changes may result in deficits and others in context-specific advantages.

The mechanisms underlying the behavioral sex-differences are not fully understood, but different concentrations of sex hormones may also play a role. For example, testosterone may facilitate cognitive task performance through decreasing response latency (Hooven et al., 2004) while estrogen activates neuroprotective signaling pathways (Arevalo et al., 2015) and is necessary for supporting spatial working memory in touchscreen tasks in females (Sbisa et al., 2017).

## Conclusions

As a complement to previous studies on MIA’s impacts, the current study reveals and characterizes additional enhancing effects of MIA on cognitive functioning. We also demonstrate that long-term EE exposure may dampen the enhancing effects of MIA when there is a loss of the enrichment in later life, providing additional support for the detrimental effects of multiple hits. Our results shed light on the variable effects of MIA in the etiology of neurodevelopmental disorders and may help facilitate potential therapeutic or preventive strategies by taking these variables into consideration.

## Supporting information

Supplementary Table S1. Maternal Immune Activation Model Reporting Guidelines Checklist

Supplementary Table S2. Prefrontal cortex mRNA expression in offspring exposed to maternal immune activation and housed in standard or environmentally

Supplementary Figure S1. Sucrose preference (%) score for male and female MIA offspring.

---

**Supplementary Figure S1.** Sucrose preference (%) score for male and female MIA offspring.

**Supplementary Table S1.** Maternal Immune Activation Model Reporting Guidelines Checklist

**Supplementary Table S2.** Prefrontal cortex mRNA expression in offspring exposed to maternal immune activation and housed in standard or environmentally enriched housing.

## Acknowledgements

This project was funded by NIMH under Award Number R15MH114035 (to ACK), a MCPHS Center for Undergraduate Research Mini-Grant (to HT), and a Summer Undergraduate Research Fellowship (to R.C.R). The authors wish to thank Drs. Johnny Kenton and Jared Young for their helpful advice on the touchscreen tasks and Dr. Steve Ramirez for his support as well. The authors would also like to thank the MCPHS University School of Pharmacy and School of Arts & Sciences for their continual support. The content is solely the responsibility of the authors and does not necessarily represent the official views of any of the financial supporters.

## Author Contributions

X.Z., H.T., H.D., & R.C.R., ran the experiments; X.Z., & A.C.K. analyzed and interpreted the data, and wrote the manuscript; A.C.K., H.T., & H.D made the figures. A.C.K. designed and supervised the study.

## Data Availability

All data are available upon request to the authors.

