## Supplementary Table S1. Maternal Immune Activation Model Reporting Guidelines Checklist for "Hidden Talents: Poly (I:C)-induced maternal immune activation improves mouse visual discrimination performance and reversal learning in a sex-dependent manner"

| ARRIVE Reporting Guideline & Recommendation | Arrive Item | MIA Model Specific Reporting Recommendation<br>***Please complete this chart for each point outlined below. If not applicable, write N/A*** |
|---|---|---|
| **Study design**<br>➢Overview of immune activation issues<br><br>For each experiment, give brief details of the study design including:<br>a. The number of experimental and control groups.<br>b. Any steps taken to minimize the effects of subjective bias when allocating animals to treatment (e.g. randomization procedure) and when assessing results (e.g. if done, describe who was blinded and when).<br>c. The experimental unit (e.g. a single animal, group or cage of animals).<br><br>A time-line diagram or flow chart can be useful to illustrate how complex study designs were carried out. | 6 | MIA Specific Reporting:<br>a. General need for improved reporting in MIA model methods + reporting pilot data<br>○ Details on pilot data:<br><br>Please see study timeline (Figure 1). |
| **Experimental procedures**<br>➢Compounds<br>➢Validation measures<br><br>For each experiment and each experimental group, including controls, provide precise details of all procedures carried out. For example:<br>a. How (e.g. drug formulation and dose, site and route of administration, anaesthesia and analgesia used [including monitoring], surgical procedure, method of euthanasia). Provide details of any specialist equipment used, including supplier(s).<br>b. When (e.g. time of day).<br>c. Where (e.g. home cage, laboratory, water maze).<br>d. Why (e.g. rationale for choice of specific anaesthetic, route of administration, drug dose used). | 7 | Provide details of:<br>a. Compounds – source, vehicle, preparation/storage, administration route, volume administered, whether anesthetics were used at time of immune challenge.<br>○ Name of compound: Polyinosine-polycytidylic acid Low molecular weight, Inv<br>○ Catalogue number: tlrl-picw<br>○ Lot number: PIW-41-03<br>○ Vehicle control used: pyrogen-free saline<br>○ Route of administration: i.p.<br>○ Volume administered: 20 mg/kg<br>○ Storage conditions: -20 degree Celsius freezer in individual aliqots<br>○ Anesthetic (type, dose, duration) used:<br>Euthanized by ketamine/xylazine (40–80 mg/kg, i.p./5–10 mg/kg, i.p.)<br>b. Housing variables at injection - temperature of room at injection time, cage change at time of injection or not<br>○ Light cycle of animal housing room: 0700-1900 Light<br>○ Time of day of injection: morning - 10am-Noon<br>○ Room temperature at injection time: RT<br>○ Did a cage change occur at time of injection: NO |

Kentner AC, Bilbo AD, Brown AS, Hsiao EY, McAllister AK, Meyer U, Pearce BD, Pletnikov MV, Yolken RH, Bauman MD. (2018). Maternal immune activation: reporting guidelines to improve the rigor, reproducibility, and transparency of the model. Neuropsychopharmacology, https://doi.org/10.1038/s41386-018-0185-7.

| | | |
|---|---|---|
| | | c. Validation of immune activation – behavior, physiological indices and/or cytokine data, including pilot dosing data<br>○ Method used to verify immune activation:<br>Validation was done by measuring body weight. Poly I:C animals either lost weight, or had a slower rate of weight gain compared to saline-treated group.<br><br>d. Validation of gestational timing – vaginal plug, estrous cycle, weight gain<br>○ Method of validating gestational timing:<br>Visualization of a copulation plug or spermatozoa in the vaginal sample after breeding. Pregnancy was confirmed by observations of a sustained diestrus phase (observed from vaginal samples), in addition to continued weight gain, and visible teats during the later phase of gestation<br><br>Additional comments:<br>N/A |
| **Experimental animals**<br>➢Species/strain/vendor<br><br>a. Provide details of the animals used, including species, strain, sex, developmental stage (e.g. mean or median age plus age range) and weight (e.g. mean or median weight plus weight range).<br>b. Provide further relevant information such as the source of animals, international strain nomenclature, genetic modification status (e.g. knock-out or transgenic), genotype, health/immune status, drug or test naïve, previous procedures, etc. | 8 | Provide details of:<br>a. Species – considerations for appropriate species (mouse, rat, non human primate, other)<br>○ Species: Mouse<br><br>b. Strain – variability in strain can influence model<br>○ Strain: C57BL/6J<br><br>c. Maternal/Offspring Physiological Variables at time of immune challenge – age, body weight<br>○ Maternal Age at challenge: gestational day 12<br>○ Maternal Body weight: ~23-26 grams<br>○ Offspring Age at challenge: offspring not re-challenged, but behavioral testinc<br>○ Offspring Sex: Both males and females tested<br>○ Offspring Body weight: ~25 grams at adult behavioral test point<br><br>d. Vendor – even within the same strain, vendor can influence endpoints<br>○ Vendor: Jackson Laboratory<br>○ Location of Vendor: Bar Harbor, Maine, USA<br>○ Room/area where animals originated from: dams came from MP15 or |

| | | |
|---|---|---|
| | | Additional Comments:<br>N/A |
| **Housing and husbandry**<br>➢Cage, ventilation, bedding, enrichment<br><br>Provide details of:<br>a. Housing (type of facility e.g. specific pathogen free [SPF]; type of cage or housing; bedding material; number of cage companions; tank shape and material etc. for fish).<br>b. Husbandry conditions (e.g. breeding program, light/dark cycle, temperature, quality of water etc for fish, type of food, access to food and water, environmental enrichment).<br>c. Welfare-related assessments and interventions that were carried out prior to, during, or after the experiment. | 9 | Provide details of:<br>a. Caging systems<br>○ *At breeding*<br>Material of cage: standard housing: polycarbonate; Enriched housir<br>Cage dimensions: standard housing: 7 1⁄2" x 11 1⁄2" x 5"; Enriched h<br>○ *After parturition*<br>Material of cage: standard housing: polycarbonate; Enriched housir<br>Cage dimensions: standard housing: 7 1⁄2" x 11 1⁄2" x 5"; Enriched h<br>○ *At weaning*<br>Material of cage: standard housing: polycarbonate; Enriched housir<br>Cage dimensions: standard housing: 7 1⁄2" x 11 1⁄2" x 5"; Enriched h<br><br>b. Animal Holding room<br>○ Temperature in room: RT (~21.Celsius)<br>○ Humidity in room: 40.6<br>○ Ventilation system: HVAC; open topped cages<br>○ Specific pathogen free [SPF]: NO<br>○ Are males & females housed in the same or separate rooms:<br>Housed in same room after weaning<br>c. Bedding exchanges/bedding type<br>○ Type of cage bedding used: Bed-o'-Cob Laboratory Animal Bedding by The /<br>○ Frequency of cage changes per week<br>*during gestation:* twice weekly<br>*during neonatal period:* no cage change until P14, then one time we<br>*following weaning:* twice weekly<br><br>d. Breeding - bred on site or timed pregnant, how many different sires (are the same fathers breeding with both experimental and control dams)<br>Breeding location: on site |

| | | |
|---|---|---|
| | | ○ Gestational age at shipping: specific age unknown, went by weight; see<br>○ Biological age of dams (if not listed in Section 8c): see above<br>○ Number of Dams bred: 51<br>○ How many times have dams been mated previously: 0<br>○ How many times did the dams mate and not become pregnant: 0-4; 7<br>○ Are the dams primiparous or multiparous? All dams are primiparous<br>○ What was the frequency of maternal handling during the gestational/neonatal period (e.g. cage cleanings, weighing, blood collection manipulations): twice weekly cage changes, daily estrous tracking<br>○ Biological age of sires: specific age unknown, went by weight (25-28 grams);<br>○ Number of sires bred: 12<br>○ How many times have sires been mated previously: mated with ~6 dams<br>○ How many times did the sires mate successfully (e.g. mating resulted in pregnancy, full term birth): almost all; 7 females did not become<br>○ If bred previously, what was the interval between mating times:<br>○ Are sires matched to experimental and control dams: YES<br>○ Describe the mating design (1:1, 1:2 etc): 1 male to 2 females<br><br>e. Social enrichment – number of cage companions<br>○ Number of cage companions prior to breeding: 2<br>○ Gestational age when dam separated for parturition: right after the<br>○ Number of cage companions at weaning: standard housed = 2-3; enriched<br><br>f. Physical enrichment – describe enrichment devices, and when enrichment is in the cage (removed when pups born? Or present throughout study), does the enrichment type change? How frequently?<br>○ Describe what type of enrichment devices (and how many) are included in cage/housing room:<br><br>Standard caging system: access to one chew bone and Nestlets® (Ancare, Bellmore, NY). Bone removed on gestational day 19 and replaced at weaning. Nestlets® available throughout study<br><br>Enrichment caging system: access to a chew bone, a tube and 1-2 toys (e.g. balls, standard pet toys, ropes, things to climb). Toys changed out twice a week. Access to toys removed on gestational day 19 and replaced at postnatal day 15. Nestlets® and Bed-r-Rest nesting disks available throughout study. |

| | | |
|---|---|---|
| | | ○ Does enrichment type/access change across study? YES<br>○ If so, when does enrichment type/access change (e.g. enrichment removed prior to parturition and replaced in late neonatal period):<br>removed on gestational day 19, prior to parturition and replaced on postnatal day 15 for enrichment animals.<br><br>Additional Comments:<br>N/A |
| **Sample size**<br>➢Litter versus offspring<br><br>a. Specify the total number of animals used in each experiment, and the number of animals in each experimental group.<br>b. Explain how the number of animals was arrived at. Provide details of any sample size calculation used.<br>c. Indicate the number of independent replications of each experiment, if relevant. | 10 | Provide details of:<br>a. Maternal N vs offspring N<br>○ What is the total number of dams/litters included in the study: 65<br>○ What is the total number of offspring per litter included the study:<br>1 male and 1 female for each test<br>b. Litter size and sex distribution<br>○ What size was each litter maintained at: 4-6 pups<br>○ What age did culling take place at: postnatal day 6 (day of birth being<br>○ How many males and females were maintained in each litter:<br>varied based on litters; tried to keep equal where possible<br>c. Cross fostering<br>○ Did cross fostering occur: NO<br>○ If so, at what age did cross fostering occur: N/A<br><br>Additional Comments:<br>N/A |

| | | |
|---|---|---|
| **Allocating animals to experimental groups**<br><br>a. Give full details of how animals were allocated to experimental groups, including randomization or matching if done.<br>b. Describe the order in which the animals in the different experimental groups were treated and assessed. | 11 | a. How many offspring per litter were used in each measure: 1 male and 1 female<br><br>b. Randomization/Matching procedures<br>○ What procedures were used to assign animals to groups:<br>pseudorandomization<br><br>c. Sex as a biological variable (behavioral and physiological outcomes)<br>○ Were both males and females evaluated in each behavioral and physiological outcome: YES<br><br>Additional Comments:<br>N/A |
| **Experimental outcomes**<br>➢ Behavioral testing<br>➢ Physiological endpoints<br><br>Clearly define the primary and secondary experimental outcomes assessed (e.g. cell death, molecular markers, behavioral changes). | 12 | a. Maternal behavior and pup interactions<br>○ If maternal care was evaluated, were there differences following immunogen challenge (if so, please briefly describe):<br><br>b. Age(s) of offspring at behavioral testing/physiological evaluation endpoints:<br>postnatal day (P) 85 physiological (collected from siblings of animals tested in touchscreeen)<br>P90 (touchscreen training)<br><br>c. Order of testing (e.g. behavioral test order)<br>○ Were animals evaluated in a counter-balanced order in terms of:<br>*presentation of tests to each animal:* NO<br>*order of experimental/control groups run through each test:* YES<br>○ What was the inter-test interval if a single animal underwent a battery of tests:<br>Animals underwent training for touchscreens and followed through with paired discrimination and [illegible] |

| | | |
|---|---|---|
| | | Additional Comments:<br>Presentation of tests were not counterbalanced because the paired discrimination test must be completed prior to the reversal learning tasks are initiated |
| **Statistical methods**<br><br>a. Provide details of the statistical methods used for each analysis.<br>b. Specify the unit of analysis for each dataset (e.g. single animal, group of animals, single neuron).<br>c. Describe any methods used to assess whether the data met the assumptions of the statistical approach. | 13 | a. Unit of analysis for each data set<br>○ Is the unit (n) of each analysis based on number of litters, or number of animals used per group:<br>n = litter |
| **Other Disclosures** | | Please make note of any other extraneous variables that you would like to report (e.g. fire alarms, construction, temporary relocations, other variables that you think we should be considering in our studies etc.):<br>N/A |

The recommended use of this reporting form is to fill it out and include it as supplemental material for each of your laboratory's research publications. If there are difficulties utilizing/adapting this fillable form, please contact one of the corresponding authors to request a copy. The authors give permission for this table to be edited for use in reporting on other animal models (e.g. postnatal immune challenge models, early life stress models) as appropriate.

Kentner AC, Bilbo AD, Brown AS, Hsiao EY, McAllister AK, Meyer U, Pearce BD, Pletnikov MV, Yolken RH, Bauman MD. (2018). Maternal immune activation: reporting guidelines to improve the rigor, reproducibility, and transparency of the model. Neuropsychopharmacology, https://doi.org/10.1038/s41386-018-0185-7.
