## Supplementary figures and images for "Hidden Talents: Poly (I:C)-induced maternal immune activation improves mouse visual discrimination performance and reversal learning in a sex-dependent manner"

### Supplementary Figure S1. Sucrose preference (%) score for male and female MIA offspring.

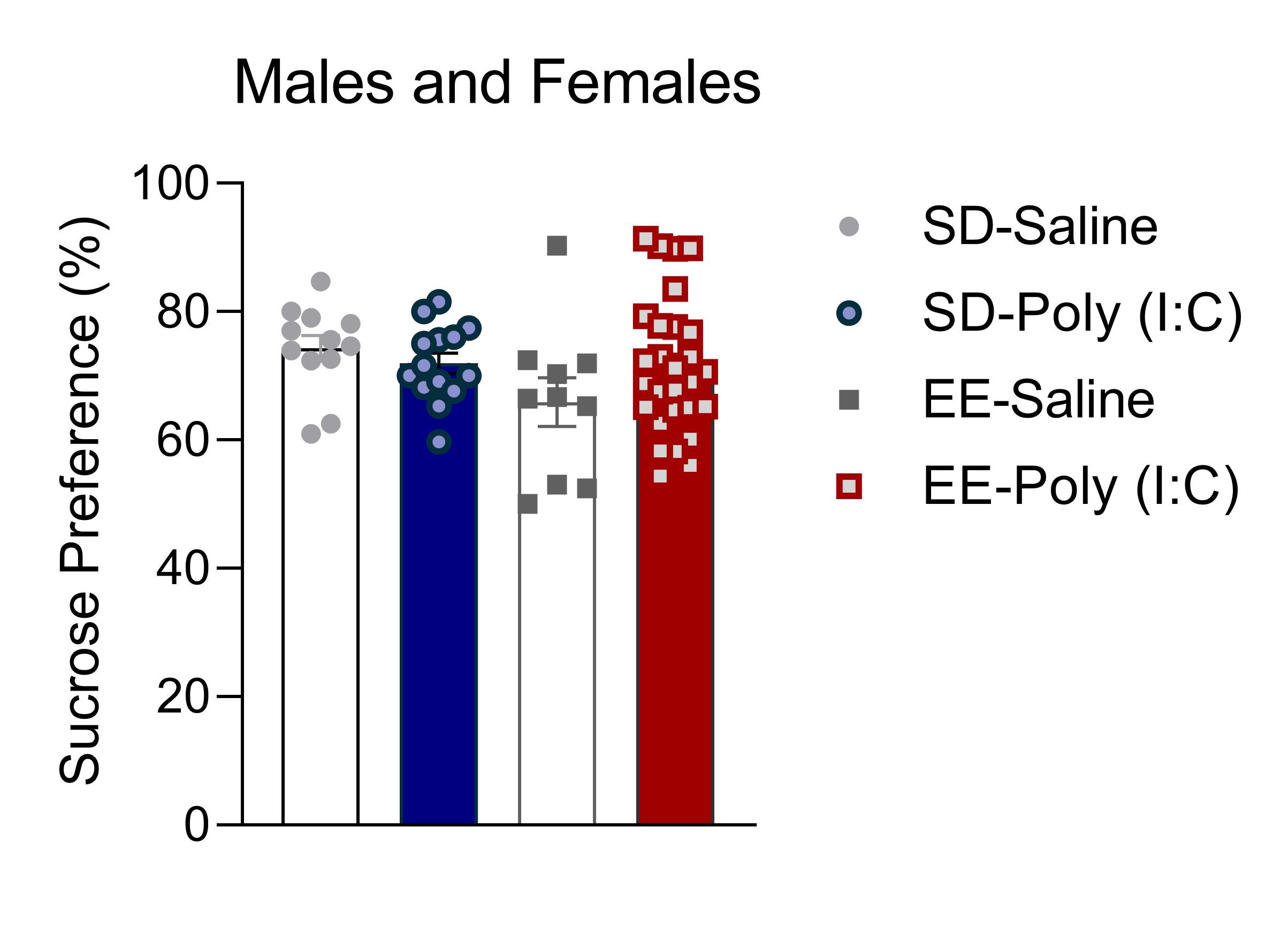

### Supplementary Table S2. Prefrontal cortex mRNA expression in offspring exposed to maternal immune activation and housed in standard or environmentally

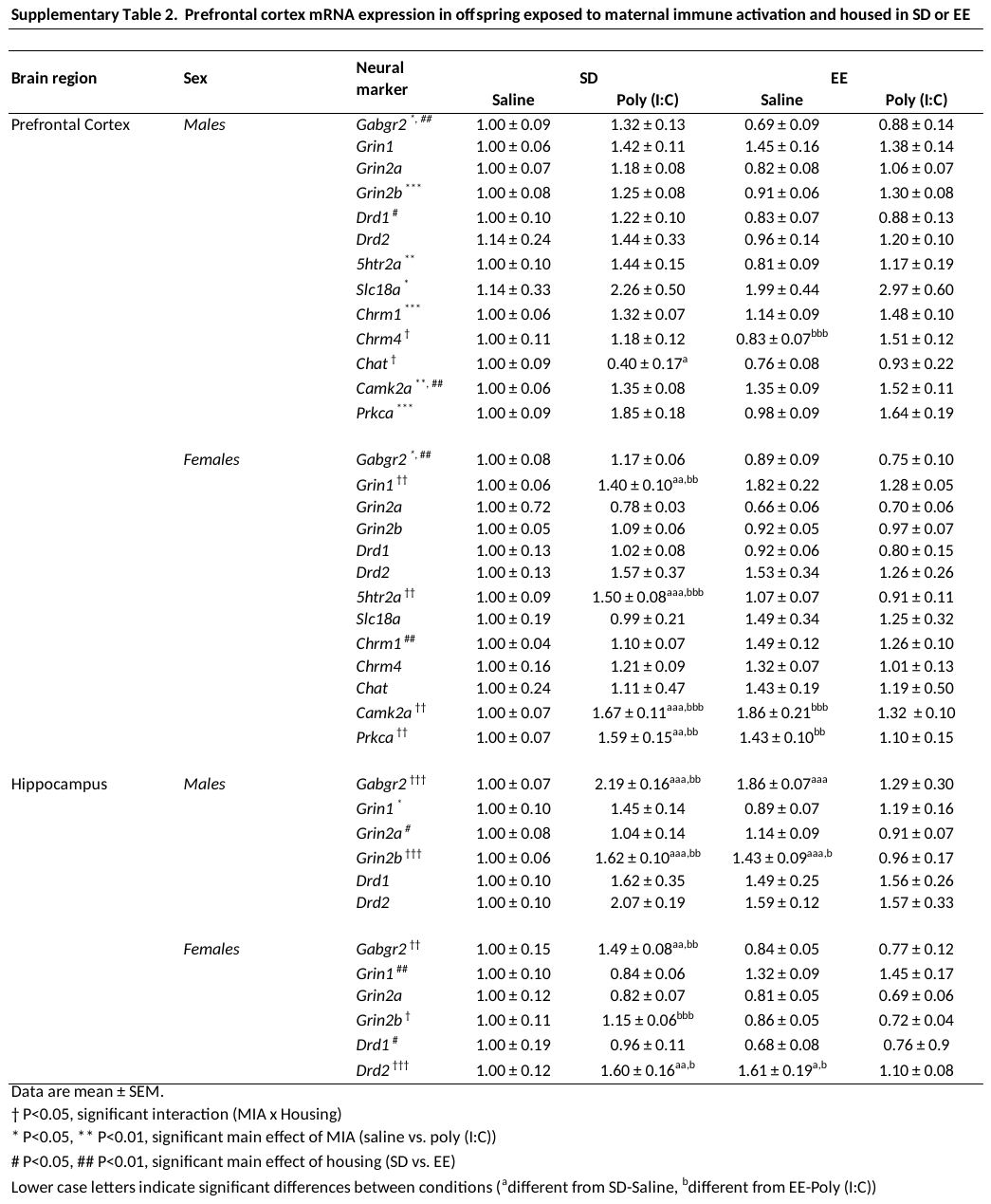
